# Climate change intensifies rapid genomic selection beyond the ancestral niche of *Fagus sylvatica*

**DOI:** 10.64898/2026.03.09.710448

**Authors:** Linda Eberhardt, Friederike Reuss, María Esther Nieto Blázquez, Jessica Hetzer, Barbara Feldmeyer, Markus Pfenninger

## Abstract

As climate change accelerates, the persistence of long-lived organisms increasingly depends on their capacity to adapt in situ. While phenotypic plasticity provides an immediate buffer, it remains uncertain whether forest trees can evolve rapidly enough to track shifting climatic niches. Here, we investigate the adaptive potential of European beech (Fagus sylvatica L.), a keystone temperate species, by leveraging different growth classes as a quasi-time-series. This approach allows us to compare growth classes established under the relatively stable climate of the early 20th century against those regenerating under contemporary warming (+1.1°C global mean temperature increase). Integrating pool-seq data from three growth classes across 43 sites in Germany with satellite-derived environmental stress indicators, we characterised past, current and projected future climate-driven selection. We detected rapid, genome-wide selective sweeps between the oldest and youngest growth classes, particularly in sites already exceeding their historical climatic niche (defined as the 95% confidence interval of pre-warming conditions). Notably, selection signatures have shifted over time: while older classes show signatures related to biotic interactions, younger cohorts exhibit intense selection on genes managing abiotic heat and drought stress. In the warmest regions, we estimated exceptionally high selection coefficients (s≈2), suggesting intense selection where beech trees exceed their ancestral niche. In older growth classes, distance and geology account for genetic differences between populations but in young growth classes climate is the primary factor, highlighting the importance of climate change. However, predictive modelling reveals a critical threshold to this resilience. While adaptive potential appears sufficient to maintain population persistence under low-emission scenarios (SSP1-2.6), high-emission trajectories (SSP5-8.5) are projected to rapidly outpace the species’ evolutionary capacity. These findings demonstrate that while trees can undergo remarkably rapid genomic shifts, the sheer velocity of unmitigated climate change threatens to exceed the fundamental limits of forest adaptation.

## 1. Introduction

Forests underpin the global carbon cycle and climate regulation (Psistaki et al., 2024). However, the increasing evolutionary pressure of climate change is rapidly altering their structure and function (Keenan, 2015). This drives niche shifts and physiological mismatches (Astigarraga et al., 2024; Illés & Móricz, 2022; Rosenblad et al., 2023) that threaten to transform forests from carbon sinks into sources (Albrich et al., 2023; Nath et al., 2024). Consequently, the persistence of these ecosystem services hinges on the capacity of forest trees to adapt to changing conditions.

For long-lived keystone species like *Fagus sylvatica* (European beech, hereafter beech), the central question is no longer whether they are affected, but whether their evolutionary machinery can keep pace with an environment changing faster than their generation time. The transition from the warmer climate of the late Pliocene to the glacial-interglacial cycles of the Pleistocene (ca. 2.6 ma ago) set off a mass extinction event of tree genera in Europe, leading to the selective filtering of those species that were able to adapt to repeated cycles of climate change through high fertility, dispersal ability and competitiveness (Kremer et al., 2025 and references therein). Some first evidence shows potential for rapid evolution of forest tree species in response to climate change due to strong gene flow, standing genetic variation and introgression, but many species are expected to lag behind this entirely unprecedented pace of warming (Alberto et al., 2013; Kremer et al., 2012; Kuparinen et al., 2010; Leites & Benito Garzón, 2023). As an anisohydric species, beech is particularly sensitive to drought as it maintains open stomata evpositen during water stress, increasing the risk of xylem embolism and hydraulic failure (Pflug et al., 2018). While introgressive hybridisation with *Fagus orientalis* has been proposed as a mechanism for beech to acquire drought- and heat-resistance alleles, the potential for such evolutionary rescue appears limited; hybrid offspring exhibit a survival rate of only 4%, likely due to reduced fitness (Stefanini et al., 2026). Given this restricted possibility for beneficial interspecific gene flow, the species’ adaptive capacity needs to rely on its existing standing diversity. The existence of such genetic variants was evidenced during the drought summers of 2018/19, which revealed highly variable responses of individual trees to water stress and highlighted that drought resilience in beech is a polygenic trait exhibiting substantial intraspecific variation (Pfenninger et al., 2021). Traditionally, high adaptive capacity of forest trees is attributed to their large effective population sizes, high gene flow, polygenic architectures, and overlapping generations acting as reservoirs for genetic diversity (Kremer et al., 2025). Nevertheless, the efficacy of this response is under scrutiny; for species such as *Fagus sylvatica*, evidence suggests that strong phenotypic plasticity may mask a risk of maladaptation through genomic offset, where complex polygenic architectures slow the pace of rapid selection (Müller et al., 2023). This concern regarding the limits of plasticity is echoed in herbaceous model systems like *Arabidopsis*. The study of alpine clines has revealed that highly plastic gene expression combined with strong purifying selection can paradoxically limit long-term adaptation and the overall rate of adaptive evolution (Hämälä et al., 2022). Taken together, these findings across both long-lived and annual plant systems suggest that while high plasticity provides a buffer against immediate environmental shifts, it may simultaneously constrain the fundamental evolutionary changes required for persistence under rapid global change. Moreover, if even short-lived herbaceous models show plasticity-constrained evolution, the risk for long-lived late-successional trees like beech is therefore likely magnified. Empirical studies are therefore urgently needed, yet studying adaptation in trees is difficult due to long generation times that prohibit experimental evolution.

While common garden experiments test potential performance in future climates, without historical samples they fall short of tracking real-time population-level genomic responses in the wild (Sigwart et al., 2025). However, long-lived organisms offer a unique opportunity to compare different generations as "living archives" in the real world, testing actual selection that has already occurred as a response to climate change *in situ*. Since selection in forest trees is strongest during early establishment (Gauzere et al., 2016), analysing successive age classes at the same location allow to observe the effects of climate change across a "quasi-time-series" experiment. Due to the long life of trees, sampling differentially aged classes allows to assess the selective effects of the warming climate of the 1980s and the current climate of 1.1°C above pre-industrial times (Bilgili et al., 2024) compared to the cooler climate of the early twentieth century. We integrate these empirical findings into predictive models to forecast the adaptive trajectory of European beech under future climate scenarios. Building on this natural experimental framework, we aimed to answer 1) to what extent forest tree populations already react to present climate change, as evidenced by genome-wide signals of selection, 2) which genes are involved in adaptation to climate change, and 3) whether observed selection signals in forest tree genomes can be linked to climate-driven environmental stress.

## 2. Material & Methods

### 2.1 Study populations

We selected 43 mature, beech-dominated stands across Germany based on climatic diversity and the presence of natural regeneration across all target growth classes (Fig. 1(a)). At each site, we sampled three distinct growth classes to represent a quasi-time-series: young growth (JU: <1 m height), pole wood (SH: 10–20 cm breast height diameter (BHD), ca. 20m tree height), and mature canopy trees (BA: >50 cm BHD) (Fig. 1(b)). Following demographic models (Genet et al., 2010), these classes correspond to tree establishment windows of 2000–2020, 1975–1995, and 1910–1930, respectively. To minimise kinship bias, sampled individuals were spaced at least 30 m apart and showed no signs of mechanical or fungal damage. At each location, 48 trees per growth class were sampled (S1). For each individual tree, 5–10 fully developed leaves were collected from lower branches, dried at 50°C for 30 minutes, and stored at -20°C until DNA extraction.

**Figure 1:**
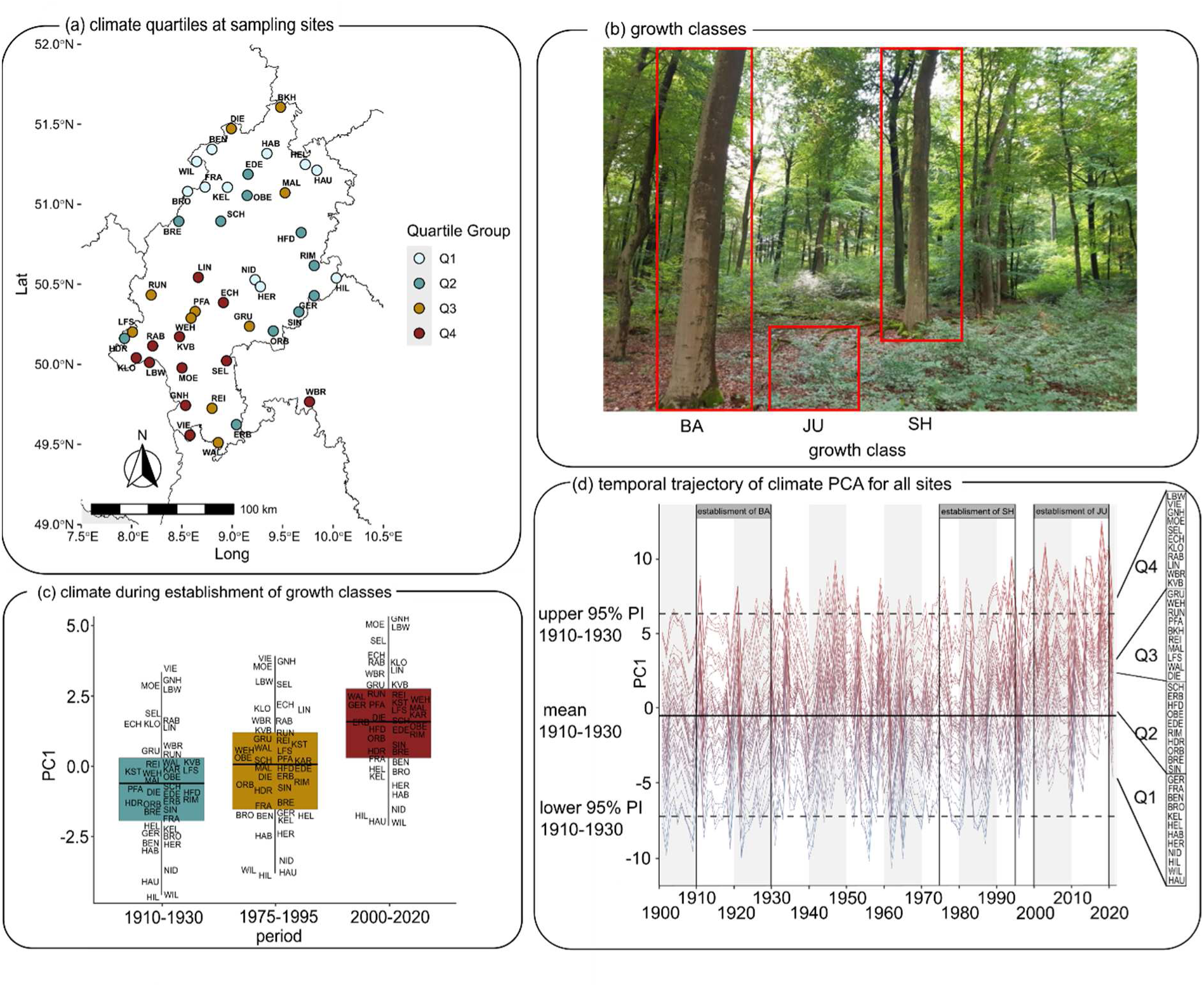
(a) Geographical distribution of sampling sites of *F. sylvatica*. Colours of points correspond to quartile groups of current climate conditions. (b) Sampled growth classes. BA: mature, canopy-forming trees of >50 cm breast-height diameter (BHD), >30 m tree height. JU: young growth of <1 m at breast height, SH: pole wood of 10-20 cm BHD, 20 m tree height. (c) Boxplot of mean PC1 scores of the site’s climate during establishment period of each growth class. (d) Temporal trajectory of climate PCA for all sampling sites from monthly grid data from 43 sampling sites for the period January 1901 to July 2021. Higher PC scores represent warmer and drier climate.

### 2.2 DNA extraction, construction of pools and sequencing

DNA was extracted from 12.5 mm² leaf disks using the NucleoMag Plant Kit (Macherey Nagel, Düren, Germany). For each site and age class, leaf samples of 48 individuals per site and growth class were pooled for DNA extraction. Sequencing libraries (450 bp insert size) were prepared by Novogene (Cambridge, UK) and sequenced on the Illumina NovaSeq 6000 platform (150 bp paired-end) to a target depth of 40X per pool. Raw sequence data are available via the European Nucleotide Archive (ENA; (Eberhardt, 2023)) under accession PRJEB64934.

### 2.3 Data processing & mapping

After initial quality control with FastQC (Andrews, 2010), reads were trimmed using Trimmomatic v0.3.9 (Bolger et al., 2014), merged with PEAR v.0.9.11 (Zhang et al., 2014), and aligned to the reference genome (Mishra et al., 2022) using BWA-MEM v.0.7.17 (Li & Durbin, 2013). Post-alignment processing, including sorting, merging, and duplicate removal, was performed using SAMtools v.1.10 (Danecek et al., 2021) and Picard v.2.20.8 (Broad Institute, 2019). We excluded repeat regions with BEDtools v.2.28.0 (Quinlan & Hall, 2010) and filtered for a minimum mean coverage of 18X per pool. Final quality metrics were generated using Qualimap v.2.2.1 (García-Alcalde et al., 2012) before converting alignments to mpileup format using SAMtools v.1.10 (Danecek et al., 2021). The toolbox Popoolation2 (Kofler, Pandey, et al., 2011) was used to convert the mpileup to sync file format and to remove indels. From the sync file, which stores raw read counts, a custom Python script was implemented to convert the data to allele frequencies relative to the reference allele. The script filters the dataset for quality and representativeness, retaining SNPs with a minimum depth of 15X per pool, a minor allele frequency (MAF) > 0.1, and < 25% missing data. To ensure marker independence and mitigate the effects of linkage disequilibrium, we subset 10,000 random positions with a minimum distance of 50 kb.

### 2.4 Climate during establishment of growth classes

#### Historical Climate and Establishment Cohorts

To characterise the climatic shifts experienced by the three tree cohorts, we retrieved monthly precipitation and temperature data (1901–2022) from the DWD (Deutscher Wetterdienst (DWD) Climate Data Center (CDC), 2025, 2025b, 2025a, 2025c). A Principal Component Analysis (PCA) was performed on scaled and centred climate variables using the R package *stats* v.3.6.2 (Posit team, 2024; R Core Team, 2024). A Bayesian paired t-test with *Bayesian First Aid* v.0.1 (Bååth, 2014) was then used to evaluate differences in mean PC1 scores across the three establishment periods: BA (1910–1930), SH (1975–1995), and JU (2000–2020). For downstream genomic analyses, sites were grouped into quartiles (Q1–Q4) based on JU-period PC1 scores, representing a gradient from cold/wet (Q1) to warm/dry (Q4) conditions.

#### Future Climate Projections and Niche Displacement

To quantify the shift of *F. sylvatica’s* climate niche beyond its ancestral niche, we projected historical and future climates using five Global Climate Models (CMIP6: GFDL-ESM4 (Dunne et al., 2020), IPSL-CM6A-LR (Boucher et al., 2020), MPI-ESM1-2-HR (Müller et al., 2018), MRI-ESM2-0 (Yukimoto et al., 2019), and UKESM1-0-LL (Sellar et al., 2019)) under three Shared Socioeconomic Pathways (SSP1-2.6, SSP3-7.0, and SSP5-8.5). Mean conditions were calculated across models for the years 2050 and 2100. Future niche displacement was quantified by projecting these scenarios onto a PCA trained on the historical baseline (1910–1930). Deviation from the ancestral niche was measured in standard units (SD) from the historical PC1 mean. Spatial analysis and mapping were performed in R using terra v.1.8-93 (Hijmans et al., 2026) and sf v.1.0-24 (Pebesma & Bivand, 2023), calculating the percentage of the study area (Hesse, Germany) projected to fall outside the historical 95% confidence interval (CI), rnaturalearth v.1.2.0 (Massicotte et al., 2023), rnaturalearthdata v.1.0.0 (South et al., 2024) and ggplot2 v.4.0.2 (Wickham, 2016).

### 2.5 Analysis of population structure and genetic variability

To assess the genetic structure across sites and cohorts, a PCA was performed on allele frequency data which were scaled, centred, and analysed via the prcomp function of the R package stats v.3.6.2. (R Core Team, 2024). Additionally, pairwise population differentiation (F_ST_) was quantified across all growth classes using PoPoolation2 (Kofler, Pandey, et al., 2011).

To detect departures from neutral equilibrium and identify genomic signatures of selection or demographic shifts, we calculated nucleotide diversity (π), Watterson’s θ, and Tajima’s D (TD) using the PoPoolation1 toolbox (Kofler, Orozco-terWengel, et al., 2011). We estimated these statistics using a sliding-window approach (1 kb windows, 1 kb step-size) with a minimum coverage of 10X and a maximum of 64X. We then employed a Bayesian implementation of a paired t-test with Bayesian First Aid v.0.2 (Bååth, 2014) to evaluate shifts in diversity and the frequency spectrum between growth classes across the climatic gradient.

### 2.6 Genome wide recombination rates

We estimated effective recombination rates to assess how selective sweeps or demographic changes influenced the genomic landscape in the oldest (BA) versus youngest (JU) cohorts, specifically comparing the most contrasting climate quartiles (Q1 and Q4). ReLERNN v.1.0.0 (Adrion et al., 2020), a deep-learning framework was used to infer recombination rates from unphased Pool-seq data.

After filtering for missingness (<20), per-base recombination rates were calculated in 10-kb non-overlapping windows across all chromosomes using ReLERNN_PREDICT (Adrion et al., 2020). We then applied ReLERNN_BSCORRECT (Adrion et al., 2020) to generate 95% confidence intervals and used Bayesian paired t-tests to identify significant differences in recombination landscapes between cohorts and climate quartiles.

### 2.7 Detection of selected sites

To identify consistent allele frequency (AF) shifts exceeding neutral expectations, Cochran-Mantel-Haenszel (CMH) tests using PoPoolation2 (Kofler, Pandey, et al., 2011) were performed. This approach leverages replicated sites within each climate quartile (Q1–Q4) to detect polygenic selection signatures while controlling for population stratification (Spitzer et al., 2020). Pairwise comparisons were conducted between growth classes within each climate quartile, applying Benjamini-Hochberg FDR correction to account for multiple testing. Significant associations were visualised via Manhattan plots using the R package qqman v. 0.1.9 (Turner, 2018).

For candidate SNPs, the selection coefficient (s) was estimated following (Lynch, 1998) and (Taus et al., 2017) to quantify the strength of selection driving AF changes between the growth classes BA and JU. To evaluate the impact of environmental stress on genomic shifts, we used a two-way ANOVA with Tukey’s HSD post-hoc tests using the R package stats v.4.4.1 to compare AF changes across the climatic gradient (Q1–Q4) and between the oldest (BA) and youngest (JU) cohorts. Effect sizes for these shifts were quantified using Cohen’s d with the package lsr v.0.5.2 (Navarro, 2015).

### 2.8 GO-term enrichment analysis of selected genes

To characterise the biological roles of the candidate loci, we identified genes associated with significant SNPs (padj<0.05) by intersecting SNP coordinates with the *F. sylvatica* genome annotation (GFF3 file). A SNP was assigned to a gene if its position fell within the genomic range (start to end) of a gene feature defined in the GFF3. Gene Ontology (GO) terms and functional domains were previously assigned to the beech genome using the InterProScan database (v. Jan. 2025) (Paysan-Lafosse et al., 2023). A GO enrichment analysis was then performed on significant SNPs for ‘Biological Processes’ using the R package topGO v.2.24.0. (Alexa & Rahnenführer, 2009) This analysis was performed on three test sets of significant SNPs from pairwise cohort comparisons (BA-SH, SH-JU, and BA-JU) within the warmest climate quartile (Q4). Overrepresented terms were identified using Fisher’s exact test with the ‘classic’ algorithm, applying a significance threshold of p<0.05.

### 2.9 Physiological Stress via Remote Sensing

Ground-level physiological stress was assessed using the Canopy Moisture Stress Index (MSI), derived from ESA Sentinel-2 Level-2 imagery (10 m resolution) (European Space Agency (ESA), 2025) via the Sentinel Hub Earth Observation Browser (Sinergise Solutions d.o.o., 2025). MSI was calculated as the ratio of shortwave infrared (SWIR, 11) to near-infrared (NIR, 8) reflectance, where values >0.54 indicate water stress in beech (Pfenninger et al., 2023). Growing season MSI (May–September, 2016–2024) was correlated with JU-period PC1 scores and genomic differentiation (Δθ between BA and JU) using Bayesian correlation tests with Bayesian First Aid (Bååth, 2014). Additionally, we employed a Hierarchical Mixed Model (*brms*) (Bürkner, 2017) and segmented regression to test if increased drought stress (higher MSI) led to higher phenotypic variability within stands, using the 0.54 threshold as a breakpoint seed.

### 2.10 Structural Equation Modelling

To partition the relative influences of geography, geology, and climate on genetic differentiation (F_ST_), we employed Path Analysis, a subset of Structural Equation Modeling (SEM) on each growth class using the R package lavaan v.0.6-21 (Rosseel, 2012). This approach allowed us to evaluate the complex, directional relationships between pairwise genetic distances and multiple environmental drivers simultaneously. We specified a model with one endogenous (response) variable, pairwise genetic distance (FST), and three exogenous (predictor) variables: geographic distance, geological distance (soil pH), and climatic distance (represented by PC1 and PC2).

To account for topographic complexity, geographic distances between stands were calculated as 3D Euclidean distances using a custom script implemented in R. Horizontal distances were determined via the Haversine formula (Sinnott, 1984) assuming an Earth radius of 6,371 km, which were then combined with altitudinal differences to provide a comprehensive metric of spatial separation. Geological variation was represented by site-specific soil pH ranges, sourced from the BGR Bodenatlas (BGR, 2025) and (Scherstjanoi, 2021). For climatic variation, we aggregated WorldClim v2.1 BioClim variables (1970–2000) (Fick & Hijmans, 2017) via PCA and calculated a Euclidean distance matrix using the axes that exceeded the broken-stick model expectation (PC1 and PC2) (available via Zenodo). All distance matrices were scaled and converted to Euclidean distances using vegan v.2.7-2 (Oksanen, 2009). The SEM was implemented using robust maximum likelihood estimation (MLR) to account for the non-independence of pairwise distance data and potential non-normality in the distance-based data. All supporting data and scripts are archived on Zenodo ((Eberhardt et al., 2026); 10.5281/zenodo.18916740).

## 3. Results

### 3.1 Rapid displacement from the ancestral climatic niche

Summer climate trajectories (May–Sept, 1901–2022) shifted significantly towards warmer and drier conditions, summarised by the first principal component (PC1; 32.9% variance). Analysis of PC1 trajectories revealed a period of relative stability until the 1980s, followed by accelerated warming and drying (Fig. 1(d)). Mean temperatures during cohort establishment rose from 7.8°C (BA) to 9.1°C (JU), a shift corroborated by Bayesian paired t-tests showing substantial climatic divergence (JU-BA: effect size = 8.0, 95% HDI [6.1,9.9], pp (β>0) = 100% (Fig. 1(c); S2). By defining the ancestral niche as the 95% confidence interval of the 1910–1930 baseline, we found that contemporary sites, particularly those in the warmest quartile (Q4), now regularly exceed the historical climatic limits at these locations. The ancestral climatic niche (1910–1930) across the study area (49°–52°N, 7.5°–10.5°E) was characterised by a range of –3.67 to 2.90 SD, with 95.7% of the landscape falling within these historical bounds. By the 1975–1995 period, the ancestral niche area began to contract, with 9.37% of the region already exceeding the upper ancestral limit. This trend accelerated in the current period (2000–2020), where the climatic range shifted upwards to 2.07–4.06 SD, leaving only 69.3% of the study area within the ancestral niche (Fig. 2(b); S3).

**Fig. 2.**
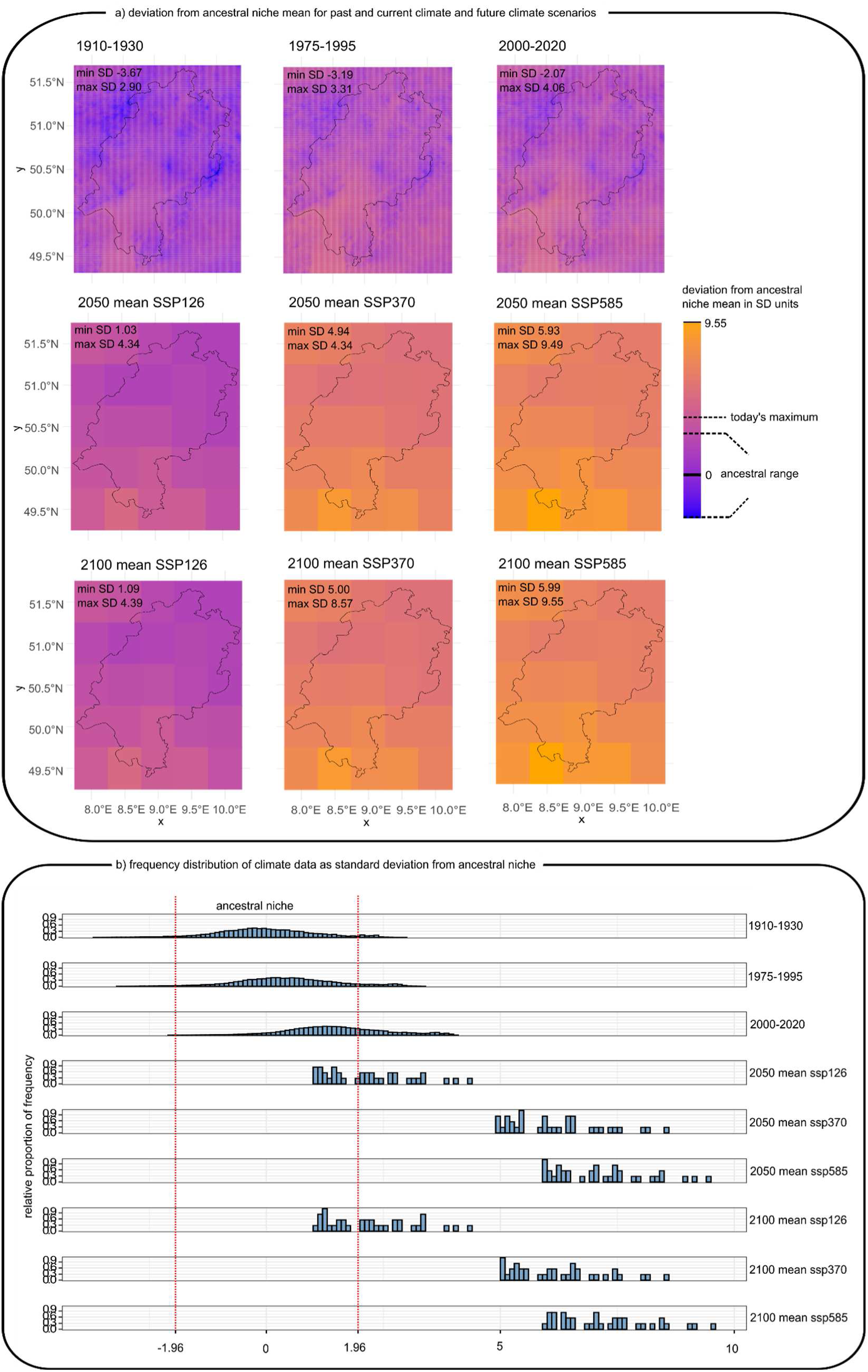
(a) Map of the spatial deviation from the ancestral niche mean (1910-1930) for past and current climate data and for three socioeconomic pathway projections for the years 2050 and 2100 in the study area. (b) Deviation from ancestral niche mean in SD units for BA, SH and JU growth classes, and for three SSP scenarios for the years 2050 and 2100 in the study area.

Future projections indicate a stark divergence based on emission pathways. Under the sustainability-focused SSP1-2.6 scenario, the climatic shift stabilises slightly; by 2100, the maximum SD remains similar to current levels, though 58.3% of the area is projected to fall outside the historical range. In contrast, under higher-emission scenarios (SSP3-7.0 and SSP5-8.5), the study area faces total climatic displacement. By 2050, the entire region is projected to shift entirely beyond the ancestral niche, with climatic deviations reaching up to 9.55 SD under SSP5-8.5 (Fig. 2(a)). These results underscore that under business-as-usual scenarios, the contemporary environment of *F. sylvatica* will become entirely novel relative to its ancestral niche, i.e. recent evolutionary history.

### 3.2 Genomic erosion and rising differentiation in the youngest cohort

#### Population Structure

After quality filtering, 114 pools (mean coverage 31X) were retained for analysis. From an initial 150 million positions, we identified 7.6 million high-quality SNPs. PCA of allele frequencies showed no discernible population structure: there were no clusters by neither growth classes from the same site, nor by climate (Fig. 3(a); PC scores: S4).

**Figure 3.**
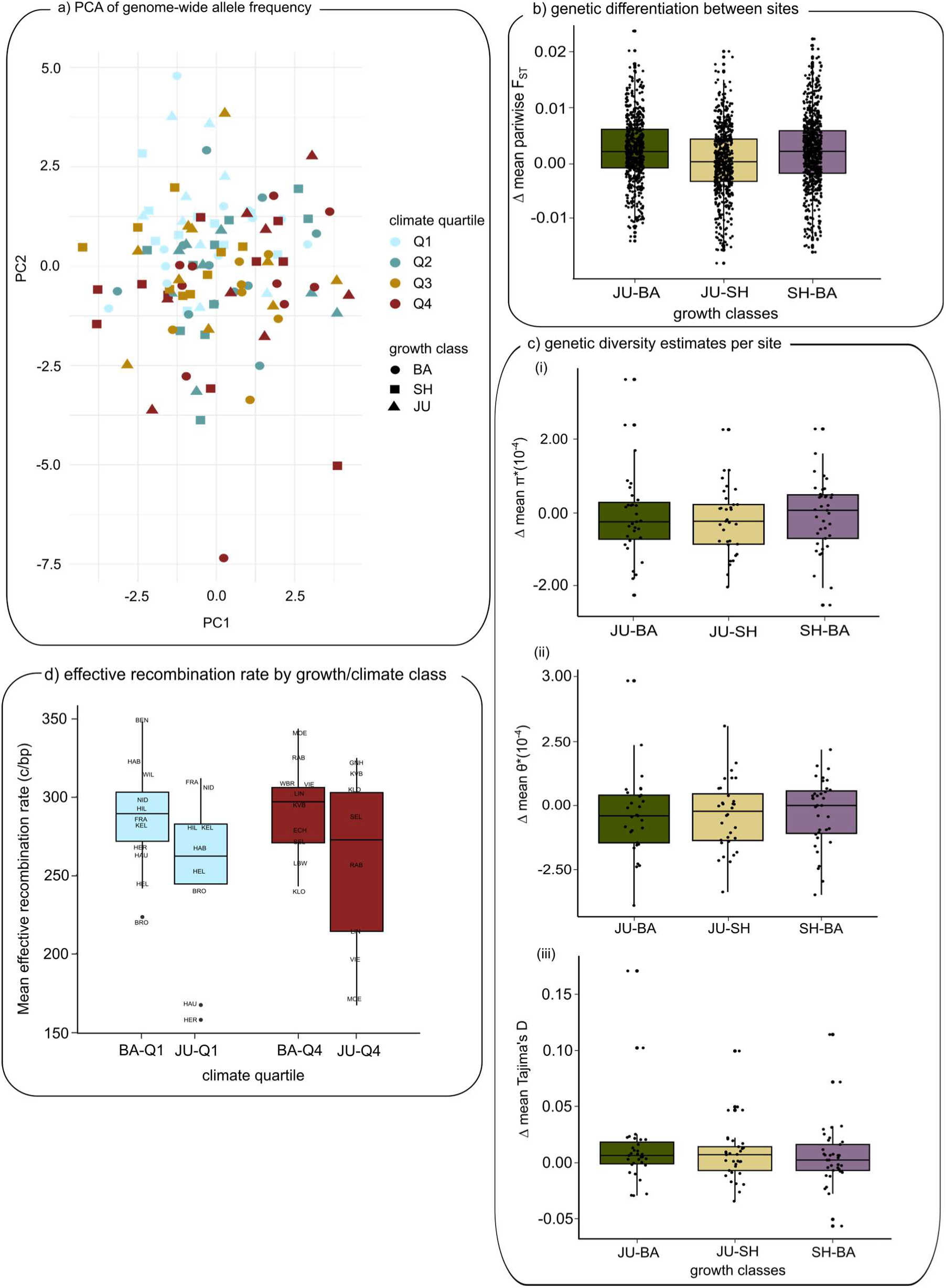
(a) PCA based on allele frequencies from three growth classes at 43 sampling sites. (b) Mean pairwise genetic differentiation (FST) between sites increased from BA to JU and from BA to SH (c(i)) Mean π per site between growth classes decreased from SH to JU. (c(ii)) Mean ϴ per site and growth class showed a decrease from BA to JU and from SH to JU. (c(iii)) Mean Tajima’s D per site (d) Mean effective recombination rate in BA and JU in the coldest quartile, Q1, and the warmest quartile, Q4, showed a decrease between BA and JU in Q4.

#### Rising Differentiation and Declining Diversity

Genetic differentiation (F_ST_) between sites increased from older to younger growth classes (JU-BA: effect size = 0.82, 95% HDI [0.70,0.92], pp (β>0) = 100%; SH-BA: effect size = 0.74, 95% HDI [0.68,0.80], pp (β>0) = 100%; JU-SH: effect size = 0.14, 95% HDI [0.073,0.20], pp (β>0) = 100%; (S5).

This rising differentiation was accompanied by an erosion of genetic diversity. Bayesian paired t-tests of both θ and π suggest decreasing trends from the older to the younger growth classes, albeit some uncertainty (Fig. 3(c)(i) and (ii), respectively) (θ(JU-BA): effect size = -0.30, 95% HDI [-0.73,0.13], pp (β<0) = 92.4%, S6; π(JU-SH): effect size = -0.25, 95% HDI [-0.64,0.11], pp (β<0) = 91.2%, S7).

Furthermore, Bayesian paired t-test of Tajima’s D between BA and JU also showed a decrease (Fig. 3(c)(iii)) (TD(JU-BA): effect size = 0.49, 95% HDI [-8.6x10-3,1.0], pp (β>0) = 97.8%; TD(SH-JU: effect size = 0.27, 95% HDI [-0.13,0.67], pp (β>0) = 90.8%; S8), suggesting that younger growth classes are deviating from neutral equilibrium.

#### Recombination Rate Decrease

Bayesian paired t-tests of effective recombination rates suggested a declining trend from BA to JU despite low posterior certainty, with the most pronounced shift occurring in the warmest sites (Fig. 3(d)) (RR(BA-JU)Q4: effect size = -0.50, 95% HDI [-1.4,0.34], pp (β<0) = 88.4%; RR(BA-JU)Q1: effect size = -0.34, 95% HDI [-1.1,0.39], pp (β<0) = 82.6% ; S9).

### 3.3 A Functional Shift of Genome-wide Selection: From Growth Optimisation to Cellular Survival in young growth class

In the warmest regions (Q4), genome-wide genetic diversity (θ) declined from BA to JU (θ(Q4(JU-BA): effect size = -0.64, 95% HDI [-1.6,0.29], pp (β<0) = 91.2%, S6), (Fig. 4(a)). Selection coefficients (s) were exceptionally high for selected alleles compared to alternative alleles (mean s = 1.89, 1.64 SD, Fig. 4(b)). In stark contrast, these same alleles conferred no selective advantage in the cooler, wetter regions (Q1–Q3; mean s = -0.04, 0.81 SD). Bayesian t-tests among Q1, Q2 and Q3 suggested an increase in selected allele frequencies, albeit with small effect sizes (all ≤ -0.34), 95% HDIs spanning [-0.6,0.014]) and some uncertainty (pp (β<0) ≤ 99.7%) (S11). In contrast, selection coefficients between Q1-Q3 and Q4 showed large effect sizes (all medians ≤ -1.2), 95% HDIs all excluding zero (spanning [-2.6,-1.4]), and maximum posterior certainty (pp (β<0) = 100%). Bayesian paired t-tests revealed markedly larger AF shifts from BA to JU involving the warmest climate quartile (Q4) compared with shifts among the cooler quartiles (Q1–Q3). Contrasts including Q4 showed huge effect sizes (all ≤ -4), 95% HDIs all excluding zero (spanning [-5,-3.5]), and very strong support ((β<0) = 100%), whereas comparisons among cooler quartiles exhibited negligible shifts and weak support (Fig. 4(c)); S12).

**Figure 4:**
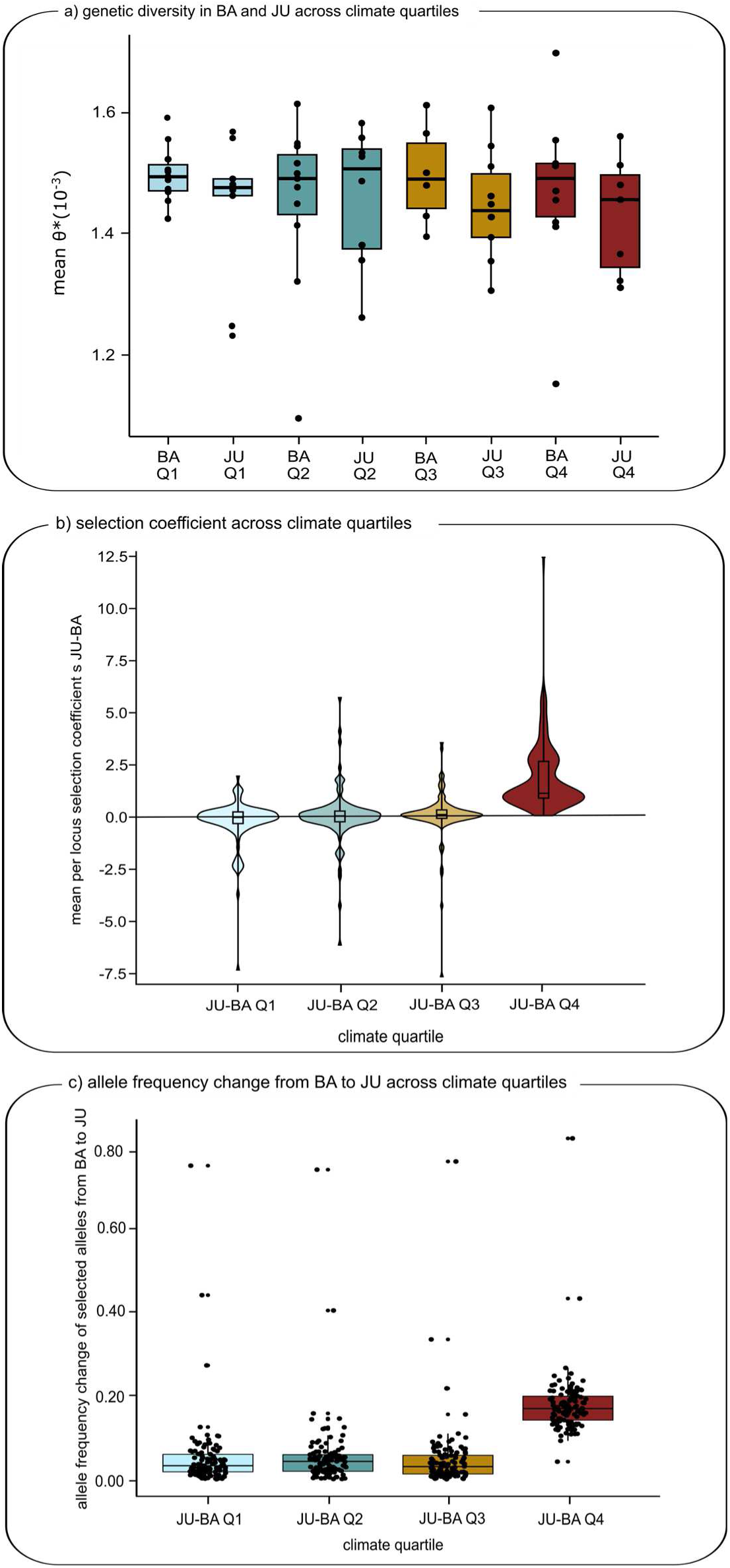
(a) Mean ϴ in BA and JU by climate quartile showed a decrease in the warmest quartile, Q4. (b) Mean per-locus selection coefficient s (Lynch, 1998; Taus et al., 2017) between BA and JU showed exceptionally high s in Q4. (c) AF change of selected SNPs between BA and JU by climate quartile showed an increase between Q1-Q2 and Q3-Q4.

#### Phenotypic Variability and Tipping Points

Hierarchical Bayesian modelling revealed that moisture stress increased the variability of phenotypic responses within stands (slope = 0.31, pp = 100%) (S15). Segmented regression identified a critical physiological breakpoint at an MSI of 0.475. Below this threshold, phenotypic variance remained stable (slope = -0.03, p = 0.28); however, beyond this inflection point, intra-stand variability increased sharply (change in slope = 0.40 per MSI unit, SE = 0.03) (S15). This suggests that once a specific drought threshold is crossed, beech stands exhibit heightened phenotypic instability, potentially signalling a loss of resilience as populations are pushed beyond their ancestral niche.

CMH tests revealed that the intensity of genomic selection tracked the climatic gradient, with the vast majority of significant allele frequency (AF) shifts occurring in the warmest quartile, with 151 loci in Q4 and 0-15 loci in Q1 to Q3 (Fig. 4(a), (b); S10). Functional enrichment analysis of the candidate loci in the warmest quartile, Q4, revealed a clear temporal transition in function of genes under selection across the three growth classes.

Selection differentiating the older growth classes (BA vs. SH) in Q4 was primarily associated with metabolic and biotic-response pathways, including sphingolipid and NADH metabolism, as well as salicylic acid-mediated signalling and innate immune responses. In contrast, selective signatures in the youngest growth class (SH vs. JU and BA vs. JU) in Q4 shifted toward cellular resilience and damage repair. Loci unique to the JU generation were significantly enriched for double-strand break repair, regulation of autophagy, telomere organization, and chromosome maintenance. This transition suggests a fundamental shift in the selective landscape: while older generations were shaped by biotic interactions and basal physiological optimisation, the youngest generation is undergoing rapid selection for cellular integrity and stress recovery in response to intensified heat and drought.

### 3.4 Environment replaces geography as major driver of population structure

#### Structural Equation Modelling: From Geography to Climate

Structural Equation Modelling (SEM) partitioned the relative effects of geography, geology, and climate on genetic differentiation (F_ST_) across the three growth classes (Fig. 6(a)(i)-(iii); S13). In the oldest growth classes (BA), genetic structure was primarily driven by geographic distance (β = 0.32, p < 0.001), consistent with a signature of isolation-by-distance (IBD). Geology exerted a significant but weaker influence (β = 0.14, p < 0.001). However, this pattern shifted fundamentally in the youngest cohort (JU), where climate dissimilarity became the dominant driver of genetic differentiation (β = 0.19, p < 0.001), superseding the relative effect of geographic distance (β = 0.17, *p* < 0.001). While the relative climatic differences between sites remained constant due to parallel shifts across the study area (Fig. 6(b), the increasing importance of climate in the JU model indicates that contemporary environmental stress is now the primary force sculpting population genetic structure.

**Fig. 5:**
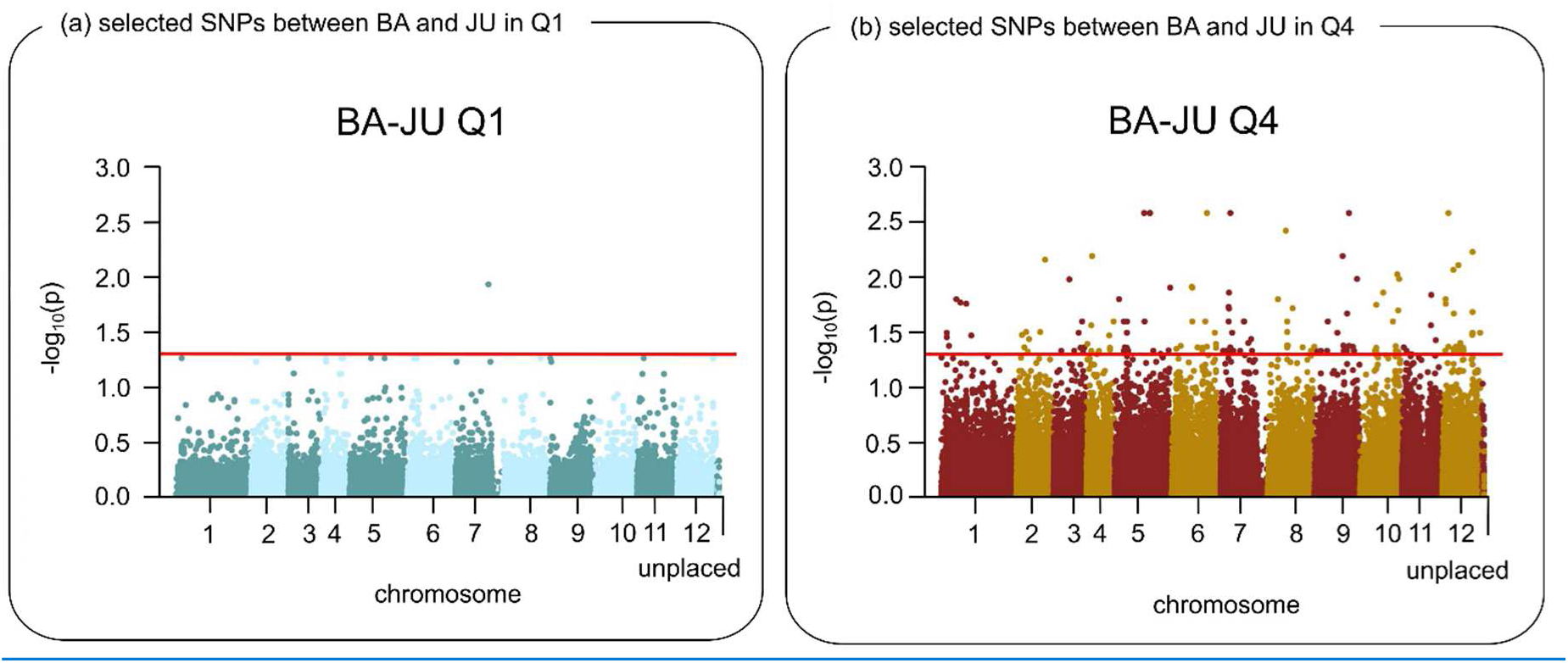
Comparison of growth classes BA and JU using CMH analysis. Manhattan plots display SNPs with Benjamini-Hochberg-adjusted p-values; one significant SNP (padj < 0.05) was detected in the coldest climate quartile, Q1 (a) and 151 in the warmest, Q4 (b).

**Figure 6:**
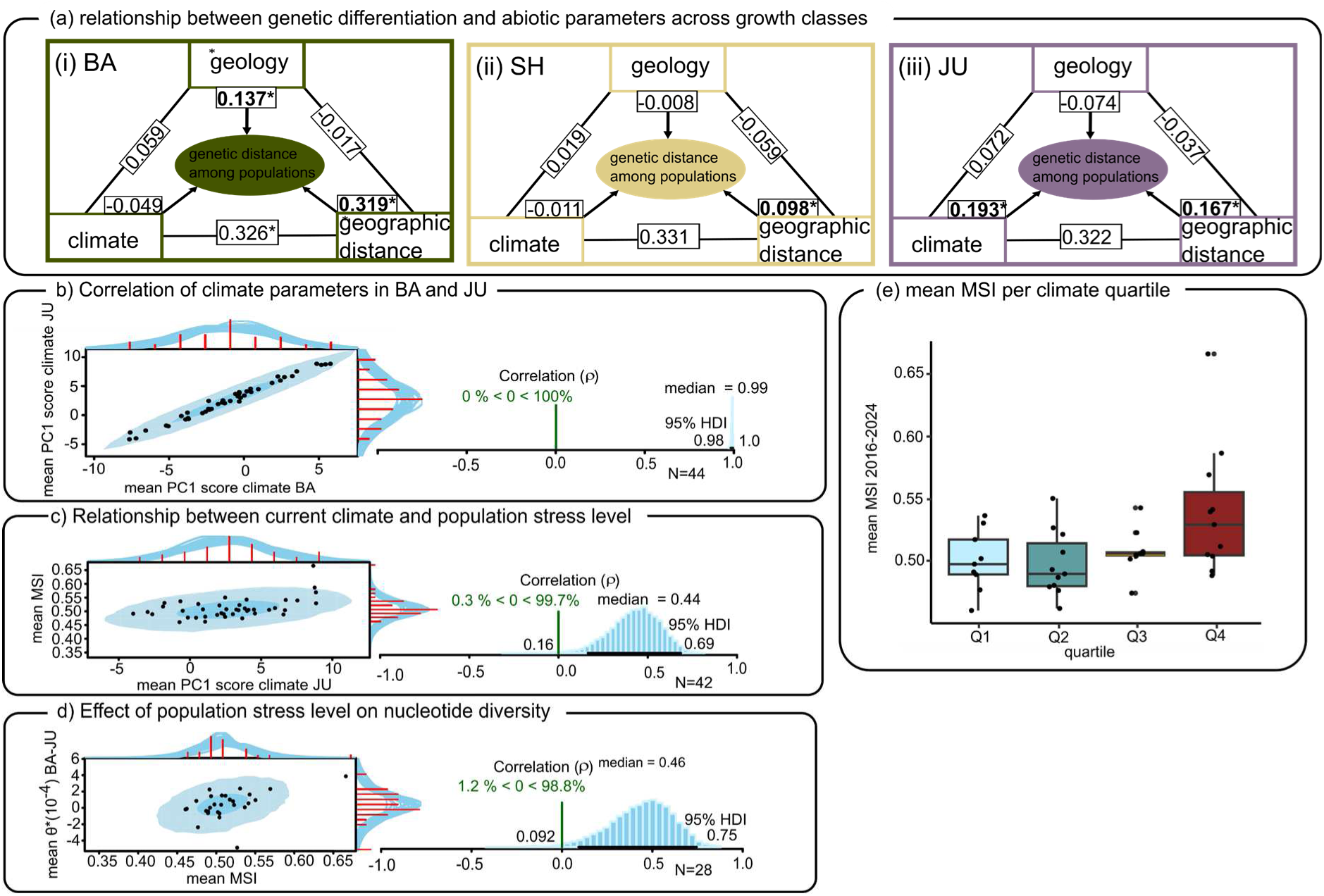
(a) Relationship diagrams based on regression coefficients of Structural equation modelling (SEM) on each growth class BA (i), SH (ii) and JU (iii) to partition the effect of geographic distance, geological variation, and climate dissimilarity on mean pairwise genetic differentiation (FST distances). (b) Climate parameters showed a strong correlation between BA and JU indicating a similar climate shift across all sites over time. (c) Correlation of climate PC1 scores and Moisture Stress Index (MSI), showing an increase in population stress levels with warmer and drier conditions. (d) Correlation of mean MSI and mean delta ϴ showed that as MSI increases, genetic diversity declines. (e) Boxplot of mean MSI by climate quartile from 2016 to 2024 showed an increase in moisture stress in the warmest quartile, Q4.

#### Linking Genomics to Physiological Stress

Satellite-derived Canopy Moisture Stress Index (MSI) validated the biological impact of these climatic shifts. We observed a strong positive correlation between JU-period climate (PC1) and mean MSI (r = 0.44, 95% HDI [0.16,0.69], pp (β>0) = 99.7%; Fig. 6(c)). Furthermore, the decline in genetic diversity between BA and JU (Δθ) was positively correlated with MSI (r = 0.46, 95% HDI [0.092,0.75], pp (β>0) = 98.8%; Fig. 6(d)), directly linking genomic erosion to moisture stress. MSI strongly increased along the climatic gradient, with the highest stress levels recorded in the warmest quartile (Q4; Fig. 6(e)).

Bayesian t-tests including Q4 showed large effect sizes (all ≤ -0.72) and strong posterior certainty ((β<0) ≥ 93.2%), despite 95% HDIs including zero (spanning [-1.9,0.22]), whereas comparisons among cooler quartiles exhibited small shifts and weak support (S14).

## 4. Discussion

### The climate experienced by established F. sylvatica cohorts changed substantially during the last 120 years

Our results demonstrate that *Fagus sylvatica* is undergoing rapid, genome-wide selection driven by anthropogenic climate warming, but this adaptive response is currently restricted to populations pushed beyond their ancestral climatic niche. Climate at the study sites remained stable from 1901 until approximately 1970, after which it shifted consistently toward warmer and drier conditions, with a marked acceleration since the mid-1980s (Fig 1(d); (Hegerl et al., 2019). We defined the ancestral niche as the 95% confidence interval of conditions during the establishment of the oldest cohort (1910–1930). Given that these conditions were representative of the preceding 200 years (Szyga, 2022), the recent climatic shift represents a significant environmental departure of past conditions. While the relative climatic positions of the 43 sites remained stable, a latitudinal and altitudinal gradient allowed us to group sites into quartiles (Q1–Q4, Fig. 1(a)). Crucially, this setup creates a natural experiment: while sites in Q1–Q3 remain largely within the ancestral niche, the youngest individuals (JU) at Q4 sites are now establishing under selective pressures previously unknown to their ancestors (Fig 1(d)).

This displacement beyond the ancestral niche allows us to test three evolutionary scenarios. First, if phenotypic plasticity is sufficient to buffer contemporary change, no systematic selection differences should be observed across any growth classes. Second, if the populations are precisely adapted to narrow local conditions, the parallel climate shift should trigger selective responses across all quartiles. Finally, if selection is driven specifically by the transgression of historical limits, we expect to find genomic signatures of adaptation exclusively in the Q4 cohort, where the environmental velocity has outpaced both plasticity and the ancestral niche.

### How far will future climate change drive F. sylvatica from its ancestral niche? Which areas will exceed the limits in the future?

To assess how different climate-change trajectories may shape the adaptive potential and viability of *F. sylvatica*, we evaluated the expected evolutionary responses under three socio-economic pathways, SSP1-2.6, SSP3-7.0 and SSP5-8.5, corresponding to low, high, and very high emissions, respectively. Under the best-case climate scenario, SSP1-2.6, the projected contraction of climate niche breadth with the same maximum deviation from the ancestral niche as observed today suggested that evolutionary adaptation to these conditions is possibly evidenced by the Q4 sites. The extremely high values even for the minimum SD projections for both the SSP3-7.0 and SSP5-8.5 scenarios represent conditions under which it is highly unlikely, if not impossible, that tree populations could adapt rapidly enough, if at all. Even with rapid sweeps, these extremes would almost certainly exceed the adaptive limits of the species, resulting in widespread tree mortality and extensive loss of beech forests.

### Population genomic analysis of genomic variation suggested little population structure and was consistent with genome-wide selective response in the JU growth class

Our results reveal a fundamental shift in the processes governing the genetic architecture of European beech. While isolation-by-distance (IBD) remains a significant driver of genetic structure across all cohorts, consistent with post-glacial recolonisation patterns (Postolache et al., 2021), the youngest generation (JU) exhibits a distinct signature of isolation-by-environment. The observed increase in F_ST_ and the concomitant decline in π and θ toward younger classes (Fig. 3(b), (c)) could theoretically result from increased drift or restricted gene flow. However, given the expanding population size of beech in Germany and the lack of significant changes in wind-mediated pollen dispersal (Bett et al., 2013), demographic shifts are unlikely. Instead, the reduction in effective recombination rates and Tajima’s D in the JU cohort points to pervasive, genome-wide selection. By linking these genomic erosions to satellite-derived moisture stress (MSI), we provide empirical evidence that climate change is currently the primary selective filter in these forests.

### Exceptional Selection at the Leading Edge

The most striking finding of this study is the magnitude of selection in the warmest climate quartile (Q4). The selection coefficients (s≈2) documented here are among the highest reported for natural populations, far exceeding the typical ranges (s < 0.5) found in meta-analyses (Thurman & Barrett, 2016). While such values might initially appear as artifacts, the consistency of the signal across independent population replicates and the high quality of the pool-seq data support their validity. These high coefficients likely reflect selection-by-mortality during the highly vulnerable seedling and sapling stages. This suggests that while beech is a long-lived species, its high fecundity and intense intraspecific competition allow for remarkably rapid adaptive tracking, in terms of generations similar to those observed in insects (Pfenninger & Foucault, 2022; Rudman et al., 2022), when environmental thresholds are crossed.

### The Shift Beyond the Ancestral Niche

Our data support a threshold-based model of adaptation. Within the ancestral niche, phenotypic plasticity appears sufficient to buffer environmental variability, maintaining the population at a neutral equilibrium. However, in the warmest quartile (Q4), the environmental velocity has outpaced the buffering capacity of plasticity, triggering intense, directional selection. This suggests that beech forests possess a "plasticity reservoir" that is currently being exhausted at the leading edge of climate change.

We identified a physiological breakpoint at an MSI of 0.475, which may serve as monitoring threshold in practical forestry. At this breakpoint, phenotypic variability rises sharply, likely reflecting the expression of cryptic genetic variation under stress (Pfenninger, Schell, et al., 2025). This shift from plastic buffering to evolutionary selection, once a characteristic of low-latitude marginal populations (Jump et al., 2006), is now emerging as a dominant process within the species’ core. As climate change progresses, the niche/edge dynamics of intense selection and reduced diversity are expanding, fundamentally altering the evolutionary trajectory of *F. sylvatica* across Europe.

### Affected genes have functions consistent with effects of climate change

The functional divergence of selective signatures between cohort comparisons (BA-SH vs. BA-JU) delineates a clear molecular timeline of *F. sylvatica*’s adaptive response to accelerating climate change. In the older BA-SH comparison, representing a period of relatively moderate climatic shifts, selection targets were primarily associated with biotic interactions and secondary metabolism.

Specifically, we observed enrichment in immunity-related pathways, salicylic acid (SA) signalling, and sphingolipid metabolism. SA is a critical phytohormone for antioxidant activation and secondary metabolite synthesis (Wani et al., 2017), but its production is often suppressed at high temperatures, leading to compromised immunity (Saijo & Loo, 2020). The simultaneous enrichment of these pathways suggest that historical selection was focused on maintaining the abiotic-biotic stress trade-off and defense against pathogens.

In contrast, the more recent BA-JU comparison reflects a shift toward pathways essential for cellular integrity under acute abiotic stress. Enrichment in cellular signalling, intracellular trafficking, and autophagy points to selection for mechanisms that sustain homeostasis under the thermal and hydric extremes of the early 21st century (Kalinowska & Isono, 2018). Notably, the enrichment of terms related to chromosome segregation and telomere organisation suggests that heat stress may already be impacting reproductive fitness. It indicates that the primary challenge for beech has shifted from "optimising growth and biotic defense" to "maintaining genomic integrity”. In *Arabidopsis*, heat stress is known to disrupt male meiosis, leading to reduced fertility (Liu et al., 2009). Our results suggest that similar heat-induced declines in reproductive performance may have acted as a powerful selective filter in beeches during the establishment of the JU cohort, potentially through compromised seed production in parent trees or reduced offspring fitness.

This transition from biotic-defence signatures to abiotic-resilience pathways indicates a fundamental change in the dominant selective filter. While older generations were primarily shaped by local biotic interactions and physiological optimisation, the youngest generation is undergoing rapid selection for survival and genomic stability as populations are pushed beyond their ancestral climatic niche. This suggests that the acceleration of climate change since the 1980s has overridden historical ecological filters, forcing a state of rapid evolutionary tracking in *F. sylvatica*.

### Increased genetic differentiation in JU class causally driven by climate change

Structural Equation Modelling (SEM) reveals a fundamental shift in the forces governing the genomic landscape of *F. sylvatica*. In the older cohorts (BA and SH), genetic differentiation was primarily driven by geography and geology (soil pH), reflecting traditional patterns of isolation-by-distance and fine-scale local adaptation to stable site conditions (Pluess et al., 2016) (Fig. 6(a)(i),(ii)). However, in the youngest growth class (JU), climatic dissimilarity emerged as the dominant driver of population differentiation (Fig. 6(a)(iii)). This transition indicates that contemporary climate selection is now superseding adaptation to historically stable, site-specific variables.

The emergence of climate as a primary driver suggests an ongoing decoupling between ancestral local adaptation and future fitness. While *F. sylvatica* has historically tracked local edaphic and spatial cues, the acceleration of climate change is imposing a novel selective filter that may lead to significant genomic load or maladaptation. As populations are pushed toward higher elevations or novel climatic states (Frank et al., 2017), they risk phenological mismatches, such as altered timing of bud break or senescence, that expose trees to late spring frosts or reduced growing seasons (Leuschner et al., 2023), at least if the substantial adaptive potential in this trade is depleted (Pfenninger, Langan, et al., 2025).

Our finding that climate did not independently predict the structure of the oldest growth class must be interpreted with caution, as the climate data used in the SEM (2000–2015) represents the environment during the youngest growth class establishment phase rather than the early 20th century. Nevertheless, the fact that the genetic structure of the youngest growth class has already aligned with this recent climate, unlike its predecessors, provides powerful evidence for rapid evolutionary tracking. This suggests that the contemporary distribution of European beech is entering a phase where ancestral adaptive signals are being overridden by the urgent necessity of surviving anthropogenic climate change.

Our results suggest that microevolutionary adaptation to contemporary climate change is occurring on a remarkably condensed timescale. While Saleh et al., (2022) demonstrated rapid adaptation in French *Quercus petraea* populations across four age cohorts spanning 340 years (1680–2008), we observed analogous genomic shifts within a mere 100-year window. Despite this significantly shorter timeframe, our findings reveal a functional convergence; both studies identified pathogen resistance and abiotic stress response as the primary targets of selection via Gene Ontology (GO) enrichment analysis.

## 5. Conclusion

This study provides empirical evidence that anthropogenic climate change is not merely a future threat but a current evolutionary force driving rapid genomic shifts in *Fagus sylvatica*. We demonstrate a clear transition in selective pressures: while older cohorts were shaped by biotic interactions and local geology, the youngest generation is being sculpted by abiotic climatic drivers, specifically heat and drought stress, as populations are forced beyond their ancestral niche. The discovery of exceptionally high selection coefficients (s≈2) in the warmest ranges confirms that long-lived forest trees possess the standing genetic variation to respond rapidly to environmental shifts. However, this adaptive capacity has limits. Our projections indicate that under high-emission scenarios (SSP3-7.0 and SSP5-8.5), the climatic displacement will far exceed the historical variance by 2100, likely outpacing even this rapid evolutionary response. Consequently, while natural selection is actively mitigating climate impacts in the short term, the long-term persistence of European beech forests hinges on stabilising climate trajectories within the physiological and evolutionary limits revealed here.

## Acknowledgements

We would like to thank Liam Langan helpful discussions on earlier drafts of the manuscript.

## Data availability statement

The data that support the findings of this study are openly available. Raw sequence data are deposited in the European Nucleotide Archive (ENA) under accession PRJEB64934 (Eberhardt, 2023). Supporting data and scripts are archived on Zenodo (Eberhardt et al., 2026) 10.5281/zenodo.18916740).

## CRediT

Linda Eberhardt: Formal Analysis, investigation, Visualization, Writing – Original Draft Preparation, Writing – Review & Editing

Friederike Reuss : Investigation, Writing – Review & Editing

María Esther Nieto Blázquez: Investigation, Software, Writing – Review & Editing

Jessica Hetzer: Investigation, Resources, Software, Writing – Review & Editing

Barbara Feldmeyer : Investigation, Resources, Writing – Review & Editing

Markus Pfenninger: Conceptualization, Formal Analysis, funding acquisition, investigation, Project Administration, Supervision, Visualization, Writing – Original Draft Preparation, Writing – Review & Editing

## References

Adrion, J., Galloway, J., & Kern, A. (2020). Predicting the Landscape of Recombination Using Deep Learning. Mol Biol Evol, 37(6), 1790–1808.

Alberto, F. J., Aitken, S. N., Alía, R., González-Marinez, S. C., Hänninen, H., Kremer, A., Lefèvre, F., Lenormand, T., Yeaman, S., Whetten, R., & Savolainen, O. (2013). Potential for evolutionary responses to climate change—Evidence from tree populations. Global Change Biology, 19(6), 1645–1661. 10.1111/gcb.12181

Albrich, K., Seidl, R., Rammer, W., & Thom, D. (2023). From sink to source: Changing climate and disturbance regimes could tip the 21st century carbon balance of an unmanaged mountain forest landscape. Forestry: An International Journal of Forest Research, 96(3), 399–409. 10.1093/forestry/cpac022

Alexa, A., & Rahnenführer, J. (2009). Gene set enrichment analysis with topGO. Bioconductor Improv, 27(1–26), 776–776.

Andrews, S. (2010). FastQC: A Quality Control Tool for High Throughput Sequence Data [Computer software]. http://www.bioinformatics.babraham.ac.uk/projects/fastqc/

Astigarraga, J., Esquivel-Muelbert, A., Ruiz-Benito, P., Rodríguez-Sánchez, F., Zavala, M. A., Vilà-Cabrera, A., Schelhaas, M.-J., Kunstler, G., Woodall, C. W., Cienciala, E., Dahlgren, J., Govaere, L., König, L. A., Lehtonen, A., Talarczyk, A., Liu, D., & Pugh, T. A. M. (2024). Relative decline in density of Northern Hemisphere tree species in warm and arid regions of their climate niches. Proceedings of the National Academy of Sciences, 121(28), e2314899121–e2314899121. 10.1073/pnas.2314899121

Bååth, R. (2014). Bayesian First Aid: A Package that Implements Bayesian Alternatives to the Classical *.test Functions in R. Proceedings of useR - the International R User Conference.

Bett, P. E., Thornton, H. E., & Clark, R. T. (2013). European wind variability over 140 yr. Advances in Science and Research, 10(1), 51–58. 10.5194/asr-10-51-2013

BGR. (2025). Bodenatlas Deutschland. Bundesanstalt Für Geowissenschaften Und Rohstoffe (BGR). https://bodenatlas.bgr.de

Bilgili, M., Tumse, S., & Nar, S. (2024). Comprehensive Overview on the Present State and Evolution of Global Warming, Climate Change, Greenhouse Gasses and Renewable Energy. Arabian Journal for Science and Engineering. 10.1007/s13369-024-09390-y

Bolger, A. M., Lohse, M., & Usadel, B. (2014). Genome analysis Trimmomatic: A flexible trimmer for Illumina sequence data. 30(15), 2114–2120. 10.1093/bioinformatics/btu170

Boucher, O., Servonnat, J., Albright, A. L., Aumont, O., Balkanski, Y., Bastrikov, V., Bekki, S., Bonnet, R., Bony, S., Bopp, L., Braconnot, P., Brockmann, P., Cadule, P., Caubel, A., Cheruy, F., Codron, F., Cozic, A., Cugnet, D., D’Andrea, F., … Vuichard, N. (2020). Presentation and Evaluation of the IPSL-CM6A-LR Climate Model. Journal of Advances in Modeling Earth Systems, 12(7), e2019MS002010. 10.1029/2019MS002010

Bürkner, P.-C. (2017). brms: An R Package for Bayesian Multilevel Models Using Stan. Journal of Statistical Software, 80(1), 1–28. 10.18637/jss.v080.i01

Danecek, P., Bonfield, J. K., Liddle, J., Marshall, J., Ohan, V., Pollard, M. O., Whitwham, A., Keane, T., McCarthy, S. A., Davies, R. M., & Li, H. (2021). Twelve years of SAMtools and BCFtools. 10(2), giab008. 10.1093/gigascience/giab008

Deutscher Wetterdienst (DWD) Climate Data Center (CDC). (2025). Grids of monthly averaged daily maximum air temperature (2m) for Germany (Version Version v19.3 & recent) [Data set]. https://opendata.dwd.de/climate_environment/CDC/

Deutscher Wetterdienst (DWD) Climate Data Center (CDC). (2025a). Grids of monthly averaged daily mean air temperature (2m) for Germany (Version Version v19.3 & recent) [Data set]. https://opendata.dwd.de/climate_environment/CDC/

Deutscher Wetterdienst (DWD) Climate Data Center (CDC). (2025b). Grids of monthly averaged daily minimum air temperature (2m) for Germany (Version Version v19.3 & recent) [Data set]. https://opendata.dwd.de/climate_environment/CDC/

Deutscher Wetterdienst (DWD) Climate Data Center (CDC). (2025c). Sum of monthly precipitation grids for Germany (Version Version v19.3 & recent) [Data set]. https://opendata.dwd.de/climate_environment/CDC/

Dunne, J. P., Horowitz, L. W., Adcroft, A. J., Ginoux, P., Held, I. M., John, J. G., Krasting, J. P., Malyshev, S., Naik, V., Paulot, F., Shevliakova, E., Stock, C. A., Zadeh, N., Balaji, V., Blanton, C., Dunne, K. A., Dupuis, C., Durachta, J., Dussin, R., … Zhao, M. (2020). The GFDL Earth System Model Version 4.1 (GFDL-ESM 4.1): Overall Coupled Model Description and Simulation Characteristics. Journal of Advances in Modeling Earth Systems, 12(11), e2019MS002015. 10.1029/2019MS002015

Eberhardt, L. (2023). Genomic basis of fitness relevant traits in F. sylvatica (No. PRJEB64934) [Data set]. European Nucleotide Archive (ENA). https://www.ebi.ac.uk/ena/browser/view/PRJEB64934

Eberhardt, L., Reuss, F., Nieto-Blázquez, M. E., Hetzer, J., Feldmeyer, B., & Pfenninger, M. (2026). *Supplementary Material for the manuscript Climate change intensifies rapid genomic selection beyond the ancestral niche of Fagus sylvatica* [Data set]. 10.5281/zenodo.18916740

European Space Agency (ESA). (2025). Modified Copernicus Sentinel data [Data set].

Fick, S., & Hijmans, R. (2017). WorldClim 2: New 1-km spatial resolution climate surfaces for global land areas. International Journal of Climatology, 37. 10.1002/joc.5086

Frank, A., Pluess, A. R., Howe, G. T., Sperisen, C., & Heiri, C. (2017). Quantitative genetic differentiation and phenotypic plasticity of European beech in a heterogeneous landscape: Indications for past climate adaptation. Perspectives in Plant Ecology, Evolution and Systematics, 26, 1–13. 10.1016/j.ppees.2017.02.001

García-Alcalde, F., Okonechnikov, K., Carbonell, J., Cruz, L. M., Götz, S., Tarazona, S., Dopazo, J., Meyer, T. F., & Conesa, A. (2012). Qualimap: Evaluating next-generation sequencing alignment data. Bioinformatics, 28(20), 2678–2679. 10.1093/bioinformatics/bts503

Gauzere, J., Klein, E. K., Brendel, O., Davi, H., & Oddou-Muratorio, S. (2016). Using partial genotyping to estimate the genetic and maternal determinants of adaptive traits in a progeny trial of Fagus sylvatica. Tree Genetics and Genomes, 12(6). 10.1007/s11295-016-1062-3

Genet, H., Bréda, N., & Dufrêne, E. (2010). Age-related variation in carbon allocation at tree and stand scales in beech (Fagus sylvatica L.) and sessile oak (Quercus petraea (Matt.) Liebl.) using a chronosequence approach. Tree Physiol., 30(2), 177–192.

Hämälä, T., Ning, W., Kuittinen, H., Aryamanesh, N., & Savolainen, O. (2022). Environmental response in gene expression and DNA methylation reveals factors influencing the adaptive potential of Arabidopsis lyrata. eLife, 11. 10.7554/eLife.83115

Hegerl, G., Brönnimann, S., Cowan, T., Friedman, A., Hawkins, E., Iles, C., Müller, W., Schurer, A., & Undorf, S. (2019). Causes of climate change over the historical record. Environmental Research Letters, 14. 10.1088/1748-9326/ab4557

Hijmans, R. J., Brown, A., & Barbosa, M. (2026). *terra: Spatial Data Analysis* (Version R package version 1.8-97) [Computer software]. https://rspatial.org/

Illés, G., & Móricz, N. (2022). Climate envelope analyses suggests significant rearrangements in the distribution ranges of Central European tree species. Annals of Forest Science, 79(1). 10.1186/s13595-022-01154-8

Institute. (2019). Picard Toolkit. https://broadinstitute.github.io/picard/

Jump, A. S., Hunt, J. M., & Peñuelas, J. (2006). Rapid climate change-related growth decline at the southern range edge of Fagus sylvatica. Global Change Biology, 12(11), 2163–2174. 10.1111/j.1365-2486.2006.01250.x

Kalinowska, K., & Isono, E. (2018). All roads lead to the vacuole—Autophagic transport as part of the endomembrane trafficking network in plants. Journal of Experimental Botany, 69(6), 1313–1324. 10.1093/jxb/erx395

Keenan, R. J. (2015). Climate change impacts and adaptation in forest management: A review. Annals of Forest Science, 72(2), 145–167. 10.1007/s13595-014-0446-5

Kofler, R., Orozco-terWengel, P., de Maio, N., Pandey, R. V., Nolte, V., Futschik, A., Kosiol, C., & Schlötterer, C. (2011). Popoolation: A toolbox for population genetic analysis of next generation sequencing data from pooled individuals. PLoS ONE, 6(1). 10.1371/journal.pone.0015925

Kofler, R., Pandey, R. V., & Schlötterer, C. (2011). PoPoolation2: Identifying differentiation between populations using sequencing of pooled DNA samples (Pool-Seq). Bioinformatics, 27(24), 3435–3436. 10.1093/bioinformatics/btr589

Kremer, A., Chen, J., & Lascoux, M. (2025). ‘Chimes of resilience’: What makes forest trees genetically resilient? New Phytologist, 246(5), 1934–1951. 10.1111/nph.70108

Kremer, A., Ronce, O., Robledo-Arnuncio, J., Guillaume, F., Bohrer, G., Nathan, R., Bridle, J., Gomulkiewicz, R., Klein, E., Ritland, K., Kuparinen, A., Gerber, S., & Schueler, S. (2012). Long-distance gene flow and adaptation of forest trees to rapid climate change. Ecology Letters, 15, 378–392. 10.1111/j.1461-0248.2012.01746.x

Kuparinen, A., Savolainen, O., & Schurr, F. (2010). Increased mortality can promote evolutionary adaptation of forest trees to climate change. Forest Ecology and Management, 259, 1003–1008. 10.1016/j.foreco.2009.12.006

Leites, L., & Benito Garzón, M. (2023). Forest tree species adaptation to climate across biomes: Building on the legacy of ecological genetics to anticipate responses to climate change. Global Change Biology, 29(17), 4711–4730. 10.1111/gcb.16711

Leuschner, C., Weithmann, G., Bat-Enerel, B., & Weigel, R. (2023). The Future of European Beech in Northern Germany—Climate Change Vulnerability and Adaptation Potential. Forests, 14(7). 10.3390/f14071448

Li, H., & Durbin, R. (2013). *Aligning sequence reads*, *clone sequences and assembly contigs with BWA-MEM* [Computer software]. https://doi.org/arXiv:1303.3997v2

Liu, Y., Xiong, Y., & Bassham, D. C. (2009). Autophagy is required for tolerance of drought and salt stress in plants. Autophagy, 5(7), 954–963. 10.4161/auto.5.7.9290

Lynch, M. W., Bruce. (1998). Genetics and analysis of quantitative traits (Vol. 1). Sinauer.

Massicotte, P., South, A., & Hufkens, K. (2023). rnaturalearth: World Map Data from Natural Earth. R Package Version 1.1.9000.

Mishra, B., Ulaszewski, B., Meger, J., Aury, J. M., Bodénès, C., Lesur-Kupin, I., Pfenninger, M., Da Silva, C., Gupta, D. K., Guichoux, E., Heer, K., Lalanne, C., Labadie, K., Opgenoorth, L., Ploch, S., Le Provost, G., Salse, J., Scotti, I., Wötzel, S., … Thines, M. (2022). A Chromosome-Level Genome Assembly of the European Beech (Fagus sylvatica) Reveals Anomalies for Organelle DNA Integration, Repeat Content and Distribution of SNPs. Frontiers in Genetics, 12(February). 10.3389/fgene.2021.691058

Müller, N. A., Geßner, C., Mader, M., Blanc-Jolivet, C., Fladung, M., & Degen, B. (2023). Genomic variation of a keystone forest tree species reveals patterns of local adaptation and future maladaptation. bioRxiv, 2023.05.11.540382-2023.05.11.540382.

Müller, W. A., Jungclaus, J. H., Mauritsen, T., Baehr, J., Bittner, M., Budich, R., Bunzel, F., Esch, M., Ghosh, R., Haak, H., Ilyina, T., Kleine, T., Kornblueh, L., Li, H., Modali, K., Notz, D., Pohlmann, H., Roeckner, E., Stemmler, I., … Marotzke, J. (2018). A Higher-resolution Version of the Max Planck Institute Earth System Model (MPI-ESM1.2-HR). Journal of Advances in Modeling Earth Systems, 10(7), 1383–1413. 10.1029/2017MS001217

Nath, D., Nath, R., & Chen, W. (2024). Faster dieback of rainforests altering tropical carbon sinks under climate change. Npj Climate and Atmospheric Science, 7(1), 235–235. 10.1038/s41612-024-00793-0

Navarro, D. (2015). Pacakge ‘lsr’.

Oksanen, J. (2009). Ordination and analysis of dissimilarities: Tutorial with R and Vegan. University Tennessee: Knoxville, TN, USA.

Paysan-Lafosse, T., Blum, M., Chuguransky, S., Grego, T., Pinto, B. L., Salazar, G. A., Bileschi, M. L., Bork, P., Bridge, A., Colwell, L., Gough, J., Haft, D. H., Letunić, I., Marchler-Bauer, A., Mi, H., Natale, D. A., Orengo, C. A., Pandurangan, A. P., Rivoire, C., … Bateman, A. (2023). InterPro in 2022. Nucleic Acids Research, 51(D1), D418–D427. 10.1093/nar/gkac993

Pebesma, E., & Bivand, R. (2023). Spatial Data Science: With applications in R. Chapman and Hall/CRC.

Pfenninger, M., & Foucault, Q. (2022). Population Genomic Time Series Data of a Natural Population Suggests Adaptive Tracking of Fluctuating Environmental Changes. Integrative and Comparative Biology, 62(6), 1812–1826. 10.1093/icb/icac098

Pfenninger, M., Langan, L., Feldmeyer, B., Eberhardt, L., Reuss, F., Hoffmann, J., Fussi, B., Seho, M., Mellert, K. H., & Hickler, T. (2025). Predicting Forest Tree Leaf Phenology Under Climate Change Using Satellite Monitoring and Population-Based Genomic Trait Association. Global Change Biology, 31(9). 10.1111/gcb.70484

Pfenninger, M., Langan, L., Feldmeyer, B., Fussi, B., & Hoffmann, J. (2023). Phenotypic drought stress prediction of European beech (Fagus sylvatica) by genomic prediction and remote sensing. 1–35.

Pfenninger, M., Reuss, F., Kiebler, A., Schönnenbeck, P., Caliendo, C., Gerber, S., Cocchiararo, B., Reuter, S., Blüthgen, N., Mody, K., Mishra, B., Bálint, M., Thines, M., & Feldmeyer, B. (2021). Genomic basis for drought resistance in european beech forests threatened by climate change. eLife, 10. 10.7554/eLife.65532

Pfenninger, M., Schell, T., Geiss, M., & Bulut, B. (2025). Pervasive and dynamic release of Cryptic Genetic Variation in Chironomus riparius: Rethinking adaptation in fluctuating environments.

Pflug, E. E., Buchmann, N., Siegwolf, R. T. W., Schaub, M., Rigling, A., & Arend, M. (2018). Resilient leaf physiological response of European beech (Fagus sylvatica L.) to summer drought and drought release. Frontiers in Plant Science, 9(February), 1–11. 10.3389/fpls.2018.00187

Pluess, A. R., Frank, A., Heiri, C., Lalagüe, H., Vendramin, G. G., & Oddou-Muratorio, S. (2016). Genome-environment association study suggests local adaptation to climate at the regional scale in Fagus sylvatica. New Phytologist, 210(2), 589–601. 10.1111/nph.13809

Posit team. (2024). RStudio: Integrated Development Environment for R (Version 2024.12.0) [Computer software]. Posit Software, PBC. http://www.posit.co/

Postolache, D., Oddou-Muratorio, S., Vajana, E., Bagnoli, F., Guichoux, E., Hampe, A., Le Provost, G., Lesur, I., Popescu, F., Scotti, I., Piotti, A., & Vendramin, G. G. (2021). Genetic signatures of divergent selection in European beech (Fagus sylvatica L.) are associated with the variation in temperature and precipitation across its distribution range. Molecular Ecology, 30(20), 5029–5047. 10.1111/mec.16115

Psistaki, K., Tsantopoulos, G., & Paschalidou, A. K. (2024). An Overview of the Role of Forests in Climate Change Mitigation. Sustainability (Switzerland), 16(14). 10.3390/su16146089

Quinlan, A. R., & Hall, I. M. (2010). BEDTools: A flexible suite of utilities for comparing genomic features. Bioinformatics, 26(6), 841–842. 10.1093/bioinformatics/btq033

R Core Team. (2024). R: A Language and Environment for Statistical Computing [Computer software]. R Foundation for Statistical Computing. https://www.R-project.org

Rosenblad, K. C., Baer, K. C., & Ackerly, D. D. (2023). Climate change, tree demography, and thermophilization in western US forests. Proceedings of the National Academy of Sciences, 120(18), e2301754120–e2301754120. 10.1073/pnas.2301754120

Rosseel, Y. (2012). lavaan: An R Package for Structural Equation Modeling. 48(2), 1–36.

Rudman, S., Greenblum, S., Rajpurohit, S., Betancourt, N., Hanna, J., Tilk, S., Yokoyama, T., Petrov, D., & Schmidt, P. (2022). Direct observation of adaptive tracking on ecological time scales in Drosophila. Science, 375. 10.1126/science.abj7484

Saijo, Y., & Loo, E. P. (2020). Plant immunity in signal integration between biotic and abiotic stress responses. New Phytologist, 225(1), 87–104. 10.1111/nph.15989

Saleh, D., Chen, J., Leplé, J.-C., Leroy, T., Truffaut, L., Dencausse, B., Lalanne, C., Labadie, K., Lesur, I., Bert, D., Lagane, F., Morneau, F., Aury, J.-M., Plomion, C., Lascoux, M., & Kremer, A. (2022). Genome-wide evolutionary response of European oaks during the Anthropocene. Evolution Letters, 6(1), 4–20. 10.1002/evl3.269

Scherstjanoi, M. (2021). pH-Werte deutscher Böden auf Wald- und Agrarflächen. 9. 10.3220/CA1632825232000

Sellar, A. A., Jones, C. G., Mulcahy, J. P., Tang, Y., Yool, A., Wiltshire, A., O’Connor, F. M., Stringer, M., Hill, R., Palmieri, J., Woodward, S., de Mora, L., Kuhlbrodt, T., Rumbold, S. T., Kelley, D. I., Ellis, R., Johnson, C. E., Walton, J., Abraham, N. L., … Zerroukat, M. (2019). UKESM1: Description and Evaluation of the U.K. Earth System Model. Journal of Advances in Modeling Earth Systems, 11(12), 4513–4558. 10.1029/2019MS001739

Sigwart, J. D., Schleuning, M., Brandt, A., Pfenninger, M., Saeedi, H., Borsch, T., Häffner, E., Lücking, R., Güntsch, A., Trischler, H., Töpfer, T., Wesche, K., & Consortium, C. (2025). Collectomics – towards a new framework to integrate museum collections to address global challenges. Natural History Collections and Museomics, 2, 1–20. 10.3897/nhcm.2.148855

Sinergise Solutions d.o.o. (2025). EO Browser. A Planet Labs company. EO Browser. https://apps.sentinel-hub.com/eo-browser/

Sinnott, R. W. (1984). Virtues of the Haversine. Sky and Telescope, 68(2), 158–158.

South, A., Michael, S., & Massicotte, P. (2024). rnaturalearthdata: World Vector Map Data from Natural Earth Used in ‘rnaturalearth’. R Package Version 0.1. 0.

Spitzer, K., Pelizzola, M., & Futschik, A. (2020). Modifying the chi-square and the cmh test for population genetic inference: Adapting to overdispersion. Annals of Applied Statistics, 14(1), 202–220. 10.1214/19-AOAS1301

Stefanini, C., Sperisen, C., Chybicki, I., Guillaume, F., Kurz, M., & Csilléry, K. (2026). Hybrid vigor and outbreeding depression after 100 years of replicate introduction of Caucasian beech to European forests. 10.22541/au.175312791.10979193/v2

Szyga, K. (2022). Assessment of Changing Agroclimatic Conditions in Poland Based on Selected Indicators. Atmosphere, 13, 1232. 10.3390/atmos13081232

Taus, T., Futschik, A., & Schlötterer, C. (2017). Quantifying Selection with Pool-Seq Time Series Data. Molecular Biology and Evolution, 34(11), 3023–3034.

Thurman, T. J., & Barrett, R. D. H. (2016). The genetic consequences of selection in natural populations. Molecular Ecology, 25(7), 1429–1448. 10.1111/mec.13559

Turner, S. (2018). qqman: An R package for visualizing GWAS results using Q-Q and manhattan plots. 10.21105/joss.00731

Wani, A. B., Chadar, H., Wani, A. H., Singh, S., & Upadhyay, N. (2017). Salicylic acid to decrease plant stress. Environmental Chemistry Letters, 15(1), 101–123.

Wickham, H. (2016). No Titleggplot2: Elegant Graphics for Data Analysis. Springer-Verlag. https://ggplot2.tidyverse.org

Yukimoto, S., Kawai, H., Koshiro, T., Oshima, N., Yoshida, K., Urakawa, S., Tsujino, H., Deushi, M., Tanaka, T., Hosaka, M., Yabu, S., Yoshimora, H., Shindo, E., Mizuta, R., Obata, A., Adachi, Y., & Ishii, M. (2019). The Meteorological Research Institute Earth System Model Version 2.0, MRI-ESM2.0: Description and Basic Evaluation of the Physical Component. Journal of the Meteorological Society of Japan. Ser. II, 97(5), 931–965. 10.2151/jmsj.2019-051

Zhang, J., Kobert, K., Flouri, T., & Stamatakis, A. (2014). PEAR: A fast and accurate Illumina Paired-End reAd mergeR. Bioinformatics, 30(5), 614–620. 10.1093/bioinformatics/btt593

